# Functional Specialization is Independent of Microstructural Variation in Cerebellum but Not in Cerebral Cortex

**DOI:** 10.1101/424176

**Authors:** Xavier Guell, Jeremy D Schmahmann, John DE Gabrieli

## Abstract

The human brain is understood to follow fundamental principles linking form (such as microstructure and anatomical connectivity) to function (perceptual, motor, cognitive, emotional, and other processes). Most of this understanding is based on knowledge of the cerebral cortex, where functional specialization is thought to be closely linked to microstructural variation as well as anatomical connectivity. The Universal Cerebellar Transform (UCT) theory has posited that the cerebellum has a different form-function organization in which microstructure is uniform, and in which functional specialization is determined solely by anatomical connectivity with extracerebellar structures. All cerebellar functions may thus be subserved by a common microstructural - and hence computational - substrate. Here we tested this hypothesis by measuring microstructural variation and functional specialization as indexed by magnetic resonance imaging in 1003 healthy humans. Cerebral cortex exhibited the expected pattern of microstructure-function correlation, but functional specialization was independent of microstructural variation in the cerebellum. These findings support the idea that cerebellar functional specialization is not determined by microstructure, and hence that cerebellar functions may be computationally constant across domains.

**SIGNIFICANCE STATEMENT:** The cerebellum is estimated to contain more than half the neurons in the human brain, is known to be involved in motoric, cognitive, and emotional functions, and is implicated in many neurological and neuropsychological disorders, but remains far less studied than the cerebral cortex. The Universal Cerebellar Transform (UCT) theory posits that one uniform computation underlies all cerebellar functions across multiple domains, but testing that idea has been difficult. Here we find that unlike the cerebral cortex, in which microstructural variation is associated with functional variation, the cerebellum exhibits relatively uniform microstructure across functionally distinct regions. These findings support UCT theory, and draw a sharp distinction between form-function relations in the cerebellum versus the cerebral cortex.

## INTRODUCTION

Anatomical, clinical, behavioral and neuroimaging evidence has determined that the cerebellum is functionally heterogenous: It is involved not only in motor control but also in multiple aspects of cognitive and affective processing (Buckner et al. 2011, Guell et al. 2015a, 2018a, Hoche et al. 2016, 2018; Kelly & Strick 2003, Middleton & Strick 1994, Schmahmann & Pandya 1991, Schmahmann 1996, Schmahmann & Pandya 1997a,b; Schmahmann & Sherman 1998, Stoodley & Schmahmann 2009). Consistent with its functional diversity, multiple studies have revealed cerebellar abnormalities in diseases that target a wide spectrum of cognitive and affective functions, ranging from Autism Spectrum Disorder (Arnold Anteraper et al. 2018) to Alzheimer’s disease (Guo et al. 2016).

How the cerebellum contributes to movement, thought, and emotion in health and disease remains an area of active debate. One line of inquiry investigates mechanisms underlying cerebellar function. Non-mutually exclusive hypotheses include *generation of internal models* (Ito 2008), *prediction* (Lesage et al. 2012), *event-timing* (Ivry & Keele 1989), *error-driven adjustment* (Ben-Yehudah et al. 2007), and *sequencing* (Molinari et al. 2008). A different line of inquiry has posited the Universal Cerebellar Transform (UCT) theory (Schmahmann 1991, 1996, 2000, 2010) that proposes that cerebellar contributions are computationally constant across domains. UCT theory is motivated by several observations: (1) cerebellar cytoarchitecture is essentially uniform (Ito 1993, Voogd & Glickstein 1998); (2) distinct regions of the cerebellum are anatomically (Kelly & Strick 2003) and functionally connected (Buckner et al. 2011) to distinct extra-cerebellar regions – in this way, UCT has access to distinct streams of information processing, and can emerge as different functions; (3) cognitive and affective symptoms after cerebellar injury show similarities with motor symptoms after cerebellar injury (Guell et al. 2015a); and (4) stimulation of two different cerebellar regions in humans results in similar functional changes in distinct regions of the cerebral cortex (Farzan et al. 2016).

Here we aim to test for the first time what is arguably the fundamental pillar of the UCT theory, namely, that microstructural variations are not associated with functional variations in the cerebellum. The absence of a relation between microstructural properties of cerebellar tissue and functional variation in the cerebellum would be a striking contrast with the cerebral cortex, in which microstructure variations are strongly related to functional variations (Amunts et al. 2007, Huntenburg et al. 2017).

To test this hypothesis we related microstructural variation and functional variation in 1003 humans from the Human Connectome Project (HPC) (Van Essen et al. 2013). The measure of microstructure was the T1w/T2w image intensity ratio, an indirect measure of myelin density (Glasser & Van Essen 2011). Myelin density is closely related to brain cytoarchitecture (Nieuwenhuys & Broere 2017). The measures of function were functional gradients that provide a low-dimensional representation of complex, high-dimensional resting-state functional connectivity patterns (Guell et al. 2018b, Margulies et al. 2016). Functional gradients thus provide a simple, continuous measure of brain function. Correlation analyses between these two measures have been able to capture microstructure-function relationships in the cerebral cortex (Huntenburg et al. 2017). Our hypothesis in the present study was that microstructural variations are not associated with functional variations in the cerebellum, and therefore that T1w/T2w intensity would not correlate with functional gradients in the cerebellum. Relating microstructure to function would provide a strong test of the idea that there are fundamentally different relations between microstructural variations and functional variations in the cerebellum versus the cerebral cortex, and the absence of a microstructure-function relation in the cerebellum would support UCT theory that cerebellum computations may be constant across functional domains.

## METHODS

### Code accessibility

All code used in this study is openly available at https://github.com/gablab/cerebellum_structurefunction

### Data

MRI data were provided by the Human Connectome Project (HCP), WU-Minn Consortium (Van Essen et al. 2013). Informed consent forms were previously approved by the Washington University Institutional Review Board. We analyzed cerebellum and cerebral cortical data from the HCP 1200 participants release, including (1) group-average resting-state connectivity data from 1003 participants as provided by HCP (four 15-minutes scans per participant; 534 were female; age was distributed as follows: 22-25 years old n=218, 26-30 n=432, 31-35 n=344, >35 n=9); (2) group-average T1w/T2w data from a subset of 881 participants as provided directly by HCP; and (3) single-participant T1w/T2w data from 100 randomly selected participants. Due to structural image quality thresholds imposed by HCP, some participants from the group average resting-state connectivity matrix (n=1003) are not included in the group average T1w/T2w data (n=881).

EPI data acquired by the WU-Minn HCP used multi-band pulse sequences (Feinberg et al. 2010, Moeller et al. 2010, Setsompop et al. 2012, Xu et al. 2013). HCP structural scans are defaced using the algorithm by (Milchenko & Marcus 2013). HCP MRI data pre-processing pipelines are primarily built using tools from FSL and FreeSurfer (Fischl 2012, Glasser et al. 2013, Jenkinson et al. 2012). HCP structural pre-processing includes cortical myelin maps generated by the methods introduced by (Glasser & Van Essen 2011). Results were visualized in surface and volumetric space using the HCP workbench view software, as well as on cerebellar flat maps using the SUIT toolbox (Diedrichsen 2006, Diedrichsen & Zotow 2015, Diedrichsen et al. 2009). Of note, cerebellum data analysis only included data from the cerebellar cortex - cerebellum deep white matter and nuclei regions are excluded in the HCP cerebellum grey matter mask.

### Experimental Design and Statistical Analysis

#### Analysis of T1w/T2w data

Cerebral cortical group-average T1w/T2w surface data from 881 participants as provided directly by HCP includes field inhomogeneity correction as described in (Glasser et al. 2013). In contrast, cerebellum data in this group-average does not include field inhomogeneity correction. We therefore conducted an additional preprocessing step of field inhomogeneity correction in volumetric data as implemented in the default bias regularization step of the SPM12 segment procedure (Ashburner & Friston 2005). This revealed a posterior (lobules Crus I/II) to anterior (lobules I-V, IX) bias (**Fig. 1A**). Given that lobules I-V are the principal motor functional regions of the cerebellum (Guell et al. 2018a, Schmahmann et al. 2009, Stoodley & Schmahmann 2009, Stoodley et al. 2012), inhomogeneity field correction is an important preprocessing step given that higher T1w/T2w intensity in lobules I-V would erroneously lead to correlations between T1w/T2w intensity and functional gradients of the cerebellum.

**Figure 1.**
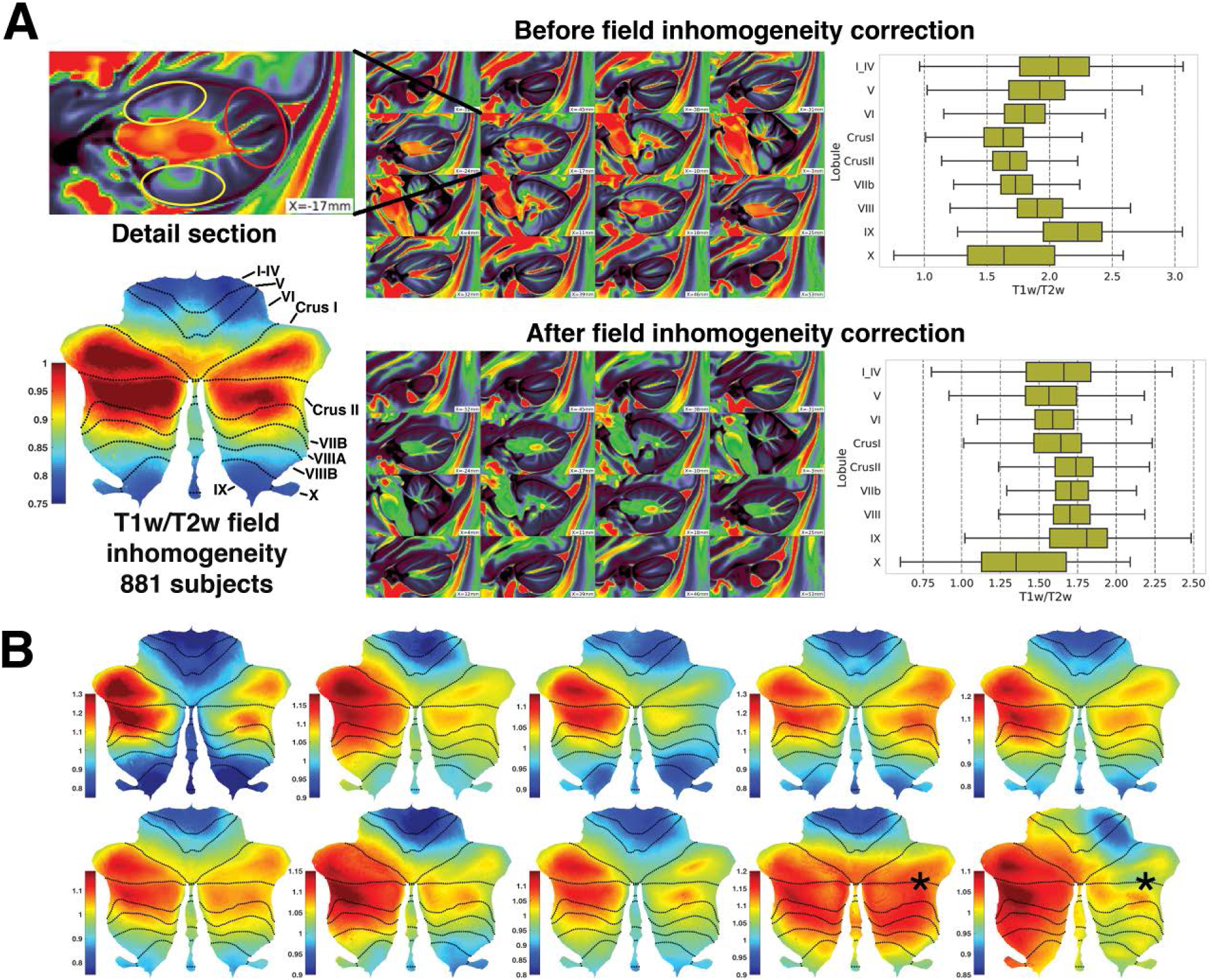
Cerebellum data corrections. **(A)**Field inhomogeneity analysis of the 881-participants group average revealed a posterior (lobules Crus I/II) to anterior (lobules I-V, IX) bias (bottom left figure in A). Before field inhomogeneity correction, higher T1w/T2w intensity was observed in the anterior aspects of the cerebellum (detail section, yellow circles) as opposed to the posterior aspects (detail section, red circles). Accordingly, lobules I-V and IX which have an anterior position in the cerebellum had higher T1w/T2w intensity before field inhomogeneity correction (top right boxplot). This effect disappeared after field inhomogeneity correction. **(B)**Field inhomogeneity analysis of 10 individual participants reveals differences in field inhomogeneity maps between individuals (for example, see region marked with an asterisk in the last two images). This raised the methodological concern that cerebellar field inhomogeneity differences may exist between participants, and hence that cerebellar field inhomogeneity correction at the average level (881 participants average) may not be optimal - supplementary analyses with field inhomogeneity correction at the single-subject level were performed to address this concern.

One methodological concern was that cerebellar field inhomogeneity differences may exist between participants, and hence that cerebellar field inhomogeneity correction at the average level (881 participants average) may not be optimal. Indeed, when analyzing field inhomogeneity in 10 individual participants, we observed between-subject differences (**Fig. 1B**). In order to address this concern, we performed cerebellar field inhomogeneity correction in a subset of 100 randomly selected participants, and averaged their T1w/T2w intensities after field inhomogeneity correction at the single-subject level. We compared the results of our analyses in the 881 participants group (with average level field inhomogeneity correction) versus in the 100 participants group (with single-subject level field inhomogeneity correction). A similar result in both cases would indicate that the analytic approach for the larger sample of 881 participants group is appropriate.

The use of T1w/T2w as a proxy measure of brain microstructure has receivied increased attention in the literature. T1w images match ex-vivo myelin histology in marmoset (Bock et al. 2009), and T1w/T2w intensity has demonstrated excellent agreement with neuroimaging maps of cytoarchitectonically defined cortical areas in humans (Glasser & Van Essen 2011, Glasser et al. 2016). Further, T1w/T2w images have very similar intensity map distributions when compared to T1 images (note that T1 sequences are different from T1w or T2w sequences in that they allow quantitative comparison across studies) (Huntenburg et al. 2017), and T1 images have been shown to correspond to ex-vivo myelin histology distribution in multiple studies (Bock et al. 2009, Geyer et al. 2011, Stüber et al. 2014). Contrasting with the small number of available T1 image datasets, T1w and T2w modalities are commonly included in very large datasets such as the HCP (n=881). Cerebral cortical microstructure-function correlations have been observed when using T1w/T2w images (Huntenburg et al. 2017), further supporting the use of T1w/T2w for the purposes of the present study.

#### Analysis of resting-state data

We calculated functional gradients in the cerebral cortex and in the cerebellum using diffusion map embedding (Coifman et al. 2005), a methodology previously used in cerebral cortical (Margulies et al. 2016) and cerebellar studies (Guell et al. 2018b). We used data from the preprocessed “dense connectome” S1200 HCP release (n=1003), which includes correlation values of each brain voxel or surface vertex with the rest of brain voxels or surface vertices. Cerebral cortical functional gradients were calculated using correlation values only within the cerebral cortex, while cerebellar functional gradients were calculated using correlation values only within the cerebellum.

Diffusion map embedding is a nonlinear dimensionality reduction technique that captures similarities between functional connectivity patterns of each voxel or surface vertex. Similar to principal component analyses, diffusion map embedding generates a first component (functional gradient 1) that accounts for as much of the variability in the data as possible. Each following gradient accounts for the highest variability possible under the constraint that all gradients are orthogonal to each other. While principal component analysis results would take the form of a mosaic (each cerebellar and cerebral cortical voxel would be assigned to a particular network with discrete borders), diffusion embedding extracts overlapping “gradients” of connectivity patterns. For example, previous cerebral cortical analyses using this method have shown that cerebral cortical functional gradient 1 extends from primary cortices to Default Mode Network (DMN) areas; gradient 2 extends from motor and auditory cortices to the visual cortex. In this way, each voxel is assigned a position within each gradient. In the preceding example, a voxel corresponding to a DMN area would be assigned an extreme position in gradient 1 (e.g. a value of 6.6 in a unitless scale from −5.5 to 6.8) and a middle position in gradient 2 (e.g. a value of 1.9 in a unitless scale from −3.1 to 5.8). The result of diffusion embedding is thus not one single mosaic of discrete networks, but multiple, continuous gradients. Each gradient reflects a given progression of connectivity patterns (e.g. from DMN to sensorimotor, from motor/auditory cortex to visual cortex, etc.), each gradient accounts for a given percentage of variability in the data, and each voxel or surface vertex has a position within each gradient.

#### T1w/T2w and functional gradients relationship analyses

Cerebral cortical resting-state and T1w/T2w maps as provided by HCP already included equal data resolution. In contrast, cerebellum T1w/T2w data had a higher resolution than cerebellum resting-state data. To allow the calculation of correlation between cerebellar T1w/T2w data and cerebellar functional gradients, cerebellar T1w/T2w data was downsampled to 2mm resolution using FSL’s FLIRT (Jenkinson & Smith 2001, Jenkinson et al. 2002). Cerebellum nifti files were then transformed to dscalar files to match the format of cerebellar functional gradient files using the -cifti-create-dense-from-template workbench command.

For cerebral cortical and cerebellar data, T1w/T2w values were plotted against functional gradient values using Seaborn’s joint histogram representation in Python. Linear, spline, and quadratic or cubic regression models were calculated using py-earth (for splines) and numpy polyfit Python tools. Py-earth is a Python implementation of Friedman’s Multivariate Adaptive Regression Splines algorithm (Friedman 1991). Splines allow the use of piecewise polynomials (for example, two linear equations) in regression analyses. Splines are thus a valuable tool when fitting nonlinear distributions - rather than using higher degree polynomials which might overfit data, piecewise polynomials such as two linear equations can offer simple models which are nonetheless able to provide high coefficients of determination (r^2^ values, i.e. the proportion of variance in one variable that is predictable from another variable in a model). Cerebellar and cerebral cortical T1w/T2w values were also grouped according to previous cerebellar (Buckner et al. 2011) and cerebral cortical parcellations (Yeo et al. 2011), and visualized in a boxplot using Seaborn’s boxplot representation in Python.

## RESULTS

### Functional gradients and T1w/T2w maps in the cerebral cortex

Functional gradients in the cerebral cortex replicated the findings by (Margulies et al. 2016) (**Fig. 2**). Gradient 1 extended from Default Mode Network (DMN) to primary cortices (sensorimotor, auditory, and visual). Gradient 2 distinguished between different primary cortical areas, extending from sensorimotor and auditory cortices to the visual cortex. Gradient 3 extended from DMN to adjacent task-positive regions. T1w/T2w revealed higher intensity values in primary cortices (motor, auditory, visual) and lower intensity values in multi-modal regions, consistent with previous reports (Glasser & Van Essen 2011, Glasser et al. 2014).

**Figure 2.**
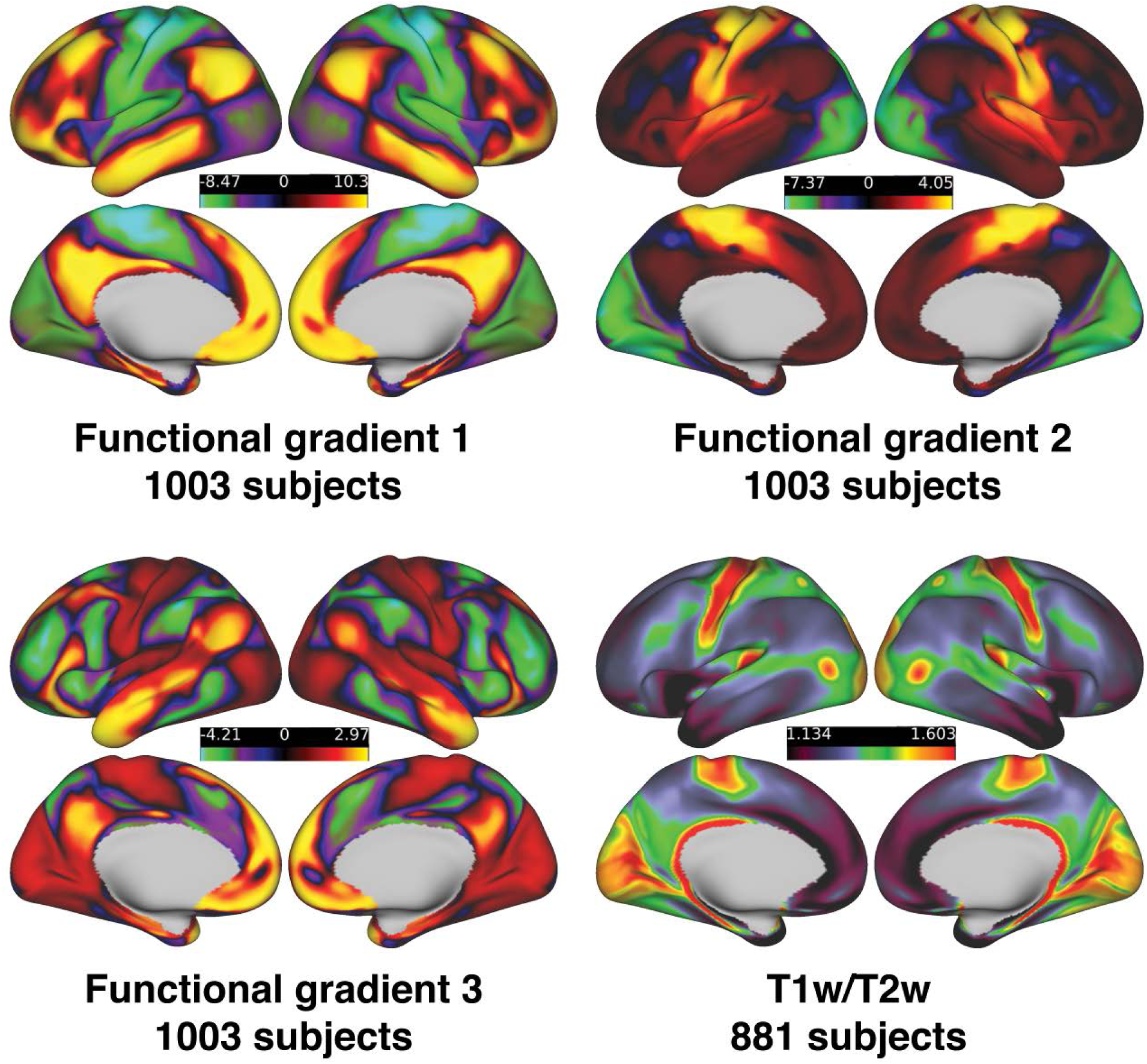
Cerebral cortical functional gradients and T1w/T2w maps.

### Functional gradients and T1w/T2w maps in the cerebellar cortex

Functional gradients in the cerebellum replicated the findings by (Guell et al. 2018b) (**Fig. 3**). Gradient 1 extended from Default Mode Network (DMN) to motor aspects of the cerebellum. Gradient 2 extended from DMN to task-positive regions. Gradient 3 distinguished between left and right aspects of the cerebellar hemispheres. T1w/T2w analysis in both the 881 average (group-level inhomogeneity correction) and 100 average (single-subject level inhomogeneity correction) analyses revealed a diffuse distribution of intensity across the cerebellar hemispheres. Higher intensity values in lobules I-IV, Crus I/II, and IX (see T1w/T2w maps in **Fig. 3**) do not correspond to any known cerebellar network distribution (Buckner et al. 2011, Guell et al. 2018a, Keren-Happuch et al. 2014, Stoodley & Schmahmann 2009, Stoodley et al. 2012).

**Figure 3.**
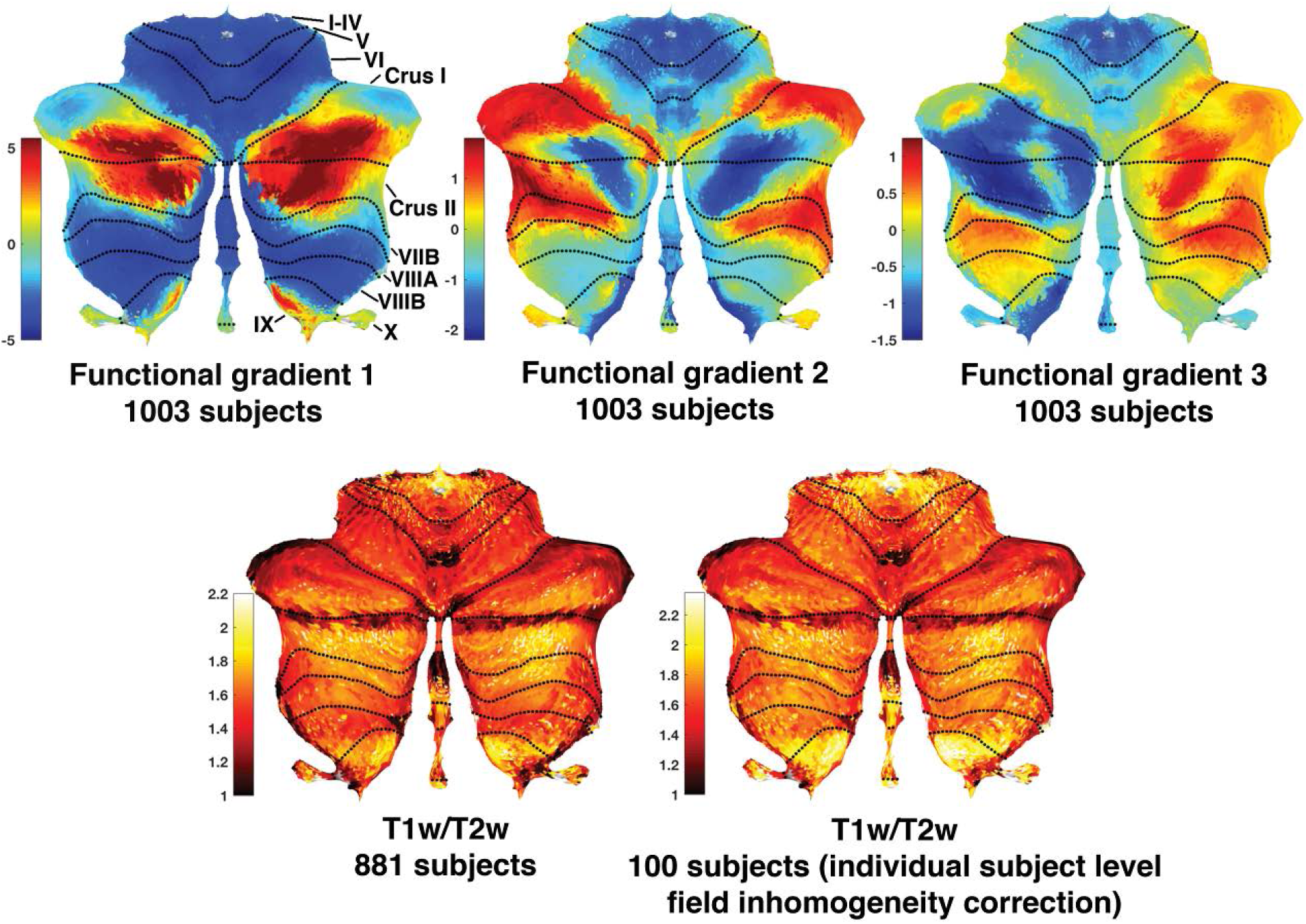
Cerebellar functional gradients and T1w/T2w maps for 881 participants group average (inhomogeneity field correction performed on average data) and 100 group average (inhomogeneity field correction performed on single-subject data before averaging).

### Relationship between functional gradients and T1w/T2w maps in the cerebellar cortex and the cerebral cortex

A strong correlation was observed between T1w/T2w values and functional gradients in the cerebral cortex, but not in the cerebellar cortex. Group average analysis (1003 participants for functional gradients, and a subset of 881 participants for T1w/T2w data) revealed r^2^ values of 0.334, 0.417, and 0.297 when fitting a model using two linear equations in gradient 1, 2, and 3, respectively. The same calculations in the cerebellum led to values of 0.025, 0.007, and 0.028. Generating a model using a higher (cubic) degree polynomial did not result in markedly improved r^2^ values. Grouping T1w/T2w values in the cerebellar cortex and the cerebral cortex based on previous cerebral cortical (Yeo et al. 2011) and cerebellar network parcellations (Buckner et al. 2011) showed little differences between networks in the cerebellum (**Fig. 4D**), contrasting with higher T1w/T2w values in primary processing networks (visual and somatomotor) in the cerebral cortex (**Fig. 4C**). While cerebellar T1w/T2w variations existed (see x axis in **Fig. 4D** showing cerebellar T1w/T2w values ranging approximately from 0.9 to 2.4), these variations were not correlated with functional gradient values, or distinct in any given network. These cerebellar T1w/T2w variations most likely correspond to variations in distance from cerebellar white matter (see volume maps in **Fig. 1A** showing higher T1w/T2w intensity in cerebellar areas close to white matter; a phenomenon that is also present in cerebral cortical T1w/T2w images). Cerebellar results did not change when using data from 100 participants who had been corrected for field inhomogeneity at the individual subject level (**Fig. 5**), supporting that failure to observe correlation between T1w/T2w intensity and functional gradients in the cerebellum is not a result of inadequate data preprocessing.

**Figure 4.**
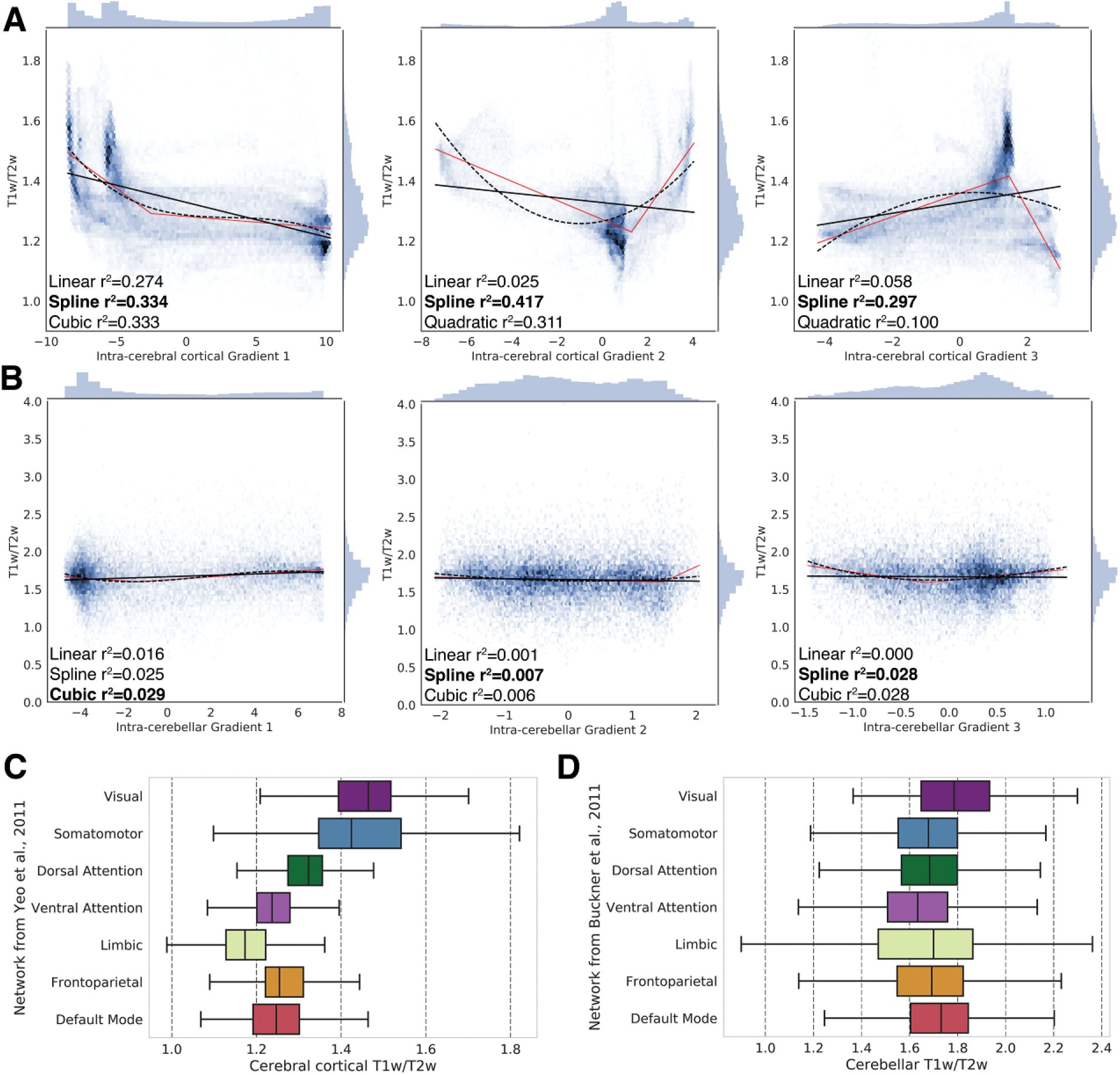
Functional gradients (1003 group average) and T1w/T2w (881 group average) relationship in the cerebellum cortex and the cerebral cortex. A strong relationship is observed between T1w/T2w values and functional gradients in the cerebral cortex **(A)**, but not in the cerebellum **(B)**. Spline regressions are shown in red lines, linear regressions are shown in black lines, quadratic or cubic regressions are shown in dotted black lines. **(C and D)** Boxes represents first to third quartile, and whiskers represent smallest and greatest datapoint within 1.5 times the interquartile range (third quartile minus first quartile). Differences of T1w/T2w intensity between networks are more pronounced in cerebral cortex **(C)** than in cerebellar cortex **(D)**. p value calculation in these analyses is not informative given the very large sample sizes (cerebellar cortex includes 17853 data points, cerebral cortex includes 59412 data points) would lead to statistically significant results even in the presence of very small effect sizes. For this reason, only data distribution visualization and effect size measures (r^2^) are presented in **(A and B)**, and only data distribution visualization is presented in **(C and D)**. Higher T1w/T2w values in cerebellar data when compared to cerebral cortical data (see x axis in **(D)**) most likely correspond to variations in distance to cerebellar white matter in the cerebellar cortex.

**Figure 5.**
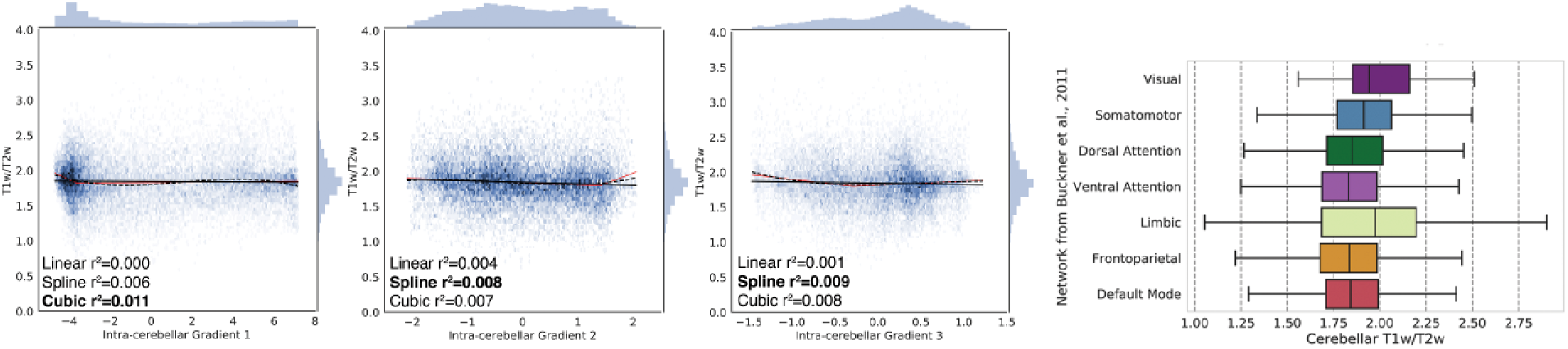
Supplementary analyses. Functional gradients (1003 group average) and T1w/T2w (100 group average, including field inhomogeneity correction at the individual-subject level before average) relationship in the cerebellar cortex. Result of this analysis is the same as in Fig. 4B, supporting that the observation of no correlation between T1w/T2w intensity and functional gradients in the cerebellum in Fig. 4B is not a result of inadequate data preprocessing.

## DISCUSSION

This study provides evidence for a fundamental dissociation in the relation between microstructure and function between the cerebellum (where there is little or no relation) versus the cerebral cortex (where there is a significant correlation). Specifically, T1w/T2w intensity (a proxy measure of microstructure) did not correlate with resting-state functional connectivity gradients (a proxy measure of function) in the cerebellum, whereas there was a strong correlation between the same measures in the cerebral cortex. These findings support the idea that cerebellar functional specialization is not determined by microstructure, and thus reinforce the UCT theory that cerebellar functions may be computationally constant across domains.

### Relevance for cerebellar neuroscience

Confirmatory evidence supporting that microstructural variations are not associated with functional variations in the human cerebellum in vivo provides new and fundamental empirical support for the Universal Cerebellar Transform theory (UCT). This theory holds that cerebellum cytoarchitecture is essentially uniform, that variations in cerebellar function arise from differences in connectivity rather than microstructure, and hence that a uniform computation underlies all cerebellar functions (Guell et al. 2017, Schmahmann 2000). Support for the UCT theory presented here advances the understanding of cerebellar function and dysfunction. First, it guides future research towards the description of an underlying cerebellar mechanism that is coherent across motor, cognitive, and affective domains. Initial hypotheses have characterized UCT as “*the cerebellar modulation of behavior, serving as an oscillation dampener maintaining function automatically around a homeostatic baseline and smoothing out performance in all domains*” (Schmahmann 2004). Second, it supports the generation of analogies between cerebellar contributions to motor, cognitive, and affective functions. For example, if the cerebellum modulates the rate, rhythm, and accuracy of movement, then it may contribute to affect by modulating the speed, capacity, consistence, and appropriateness of emotions (Guell et al. 2017, Schmahmann 1991). Third, it informs the understanding of cognitive and affective symptoms in diseases that affect the cerebellum. As the UCT theory states that a uniform mechanism underlies all cerebellar functions, nonmotor symptoms in cerebellar dysfunction may be the manifestation of the same neurological impairment that underlies cerebellar motor ataxia (“Dysmetria of Thought” theory) (Schmahmann 1991, 1996, 2000, 2010).

### Relevance for cerebral cortical neuroscience

A previous study described a strong and significant correlation between T1w/T2w intensity and cerebral cortical functional gradient 1 (see supplementary material in (Huntenburg et al. 2017)). Our analysis replicates this finding. Further, our study is the first to reveal a correlation between T1w/T2w intensity and cerebral cortical functional gradients 2 and 3.

Gradient 1 extends from Default Mode Network (DMN) to primary cortical areas (sensorimotor, auditory, and visual) (Margulies et al. 2016). Previous studies have shown higher myelin content in primary sensorimotor, visual, and auditory cortices, and lower myelin content in multimodal association areas (Glasser & Van Essen 2011, Huntenburg et al. 2017). Myelination is a crucial and precise modulator of axon conduction speed (Salami et al. 2003). At the same time, myelin consolidates neural circuits, reducing plasticity (McGee et al. 2005). In this way, myelin density differences between primary and transmodal areas might have evolved to enable not only higher conduction speed in primary cortices, but also higher plasticity in transmodal regions (Glasser & Van Essen 2011, Huntenburg et al. 2017). The linear relationship observed here between T1w/T2w intensity and cerebral cortical functional gradient 1 replicates Huntenburg’s observation that microstructural variations are associated with functional variations in the cerebral cortex.

The present findings extend this observation by showing that a strong and statistically significant microstructure-function correlation also exists in cerebral cortical functional gradients 2 and 3, with the caveat that this relationship is not linear. Gradient 2 separates distinct primary cortical areas, extending from sensorimotor and auditory to visual cortex. Primary cortical areas are thus located at both extreme poles of gradient 2, with transmodal (Default Mode Network) regions located at a middle position. The regression analysis using two linear equations was thus able to capture a strong and significant “V shape” relationship, with a coefficient of determination even higher than that observed in gradient 1. Gradient 3 distinguishes between task-positive and task-negative (DMN) networks, situating primary cortices at a middle position. Hence, two linear equations in our analysis were also able to capture a strong and significant “inverted V shape” correlation. These findings further reinforce the notion that microstructure relates to function in the cerebral cortex, not only as captured by gradient 1 (primary to transmodal regions), but also as captured by additional orthogonal gradients of functional organization.

### Limitations

As T1w/T2w ratio is not independent of MRI sequence parameters, between-subject comparisons require additional corrections. Given that we did not perform any between-subject T1w/T2w statistical comparisons other than averaging, this limitation is not immediately relevant in the present study. Further corrections such as using relative percent difference values (as suggested by (Glasser & Van Essen 2011)) or using additional calibration algorithms (Ganzetti et al. 2014) were therefore not performed in the present study beyond what was already implemented in HCP.

Cerebellar field inhomogeneity correction was performed at the group average level, a potentially suboptimal approach given that field inhomogeneity might differ between participants. Indeed, field inhomogeneity analysis in a subsample of 10 participants revealed some differences between subjects (**Fig. 1B**). We therefore calculated cerebellar field inhomogeneity correction at the individual subject level in a subset of 100 participants. Analysis in this subsample of 100 participants revealed very similar results to our initial calculations where field inhomogeneity correction was performed at the average level (note the similarity between the results in **Fig. 4B** and **Fig. 5**). This observation provides reassurance that the lack of correlation between cerebellar T1w/T2w intensity and functional gradients in our larger analysis of 881 participants was not a result of inadequate data preprocessing.

Previous investigations have highlighted the presence of anatomical and physiological variation across the cerebellar cortex. Some authors interpret these characteristics as proof against the concept of a Universal Cerebellar Transform (Cerminara et al. 2015). However, as we have argued previously (Guell et al. 2017, Schmahmann 2000), the UCT theory is based on the notion of an essentially (not completely) uniform cytoarchitecture. Uniformity far exceeds the presence of minor variations across the cerebellar cortex, in clear contrast with major microstructural variations that are observed across the cerebral cortex.

### Future studies

The present study employed T1w/T2w intensity and functional gradients to evaluate the relation of microstructural variation to functional variation in the cerebellum. Future studies may investigate other measures of topographical variation. For example, gene expression studies have revealed sagitally-oriented aggregates of distinct transcriptional patterns (“zebrin stripes”) in the cerebellum (Oberdick et al. 1998). T1w/T2w intensity has been correlated with transcriptional variation in the human cerebral cortex (Burt et al. 2018), and this motivates further research as to whether a similar or distinct relationship between transcriptional variation and microstructure or function can be observed in the cerebellum. Whereas functional gradients or T1w/T2w in the cerebellum do not seem to obey a parasagittal organization (**Fig. 3**), fMRI task activation maps in multiple motor and nonmotor domains might follow a somewhat parasagittal distribution (see figure 2 and supplementary figure 1 in (Guell et al. 2018a). Further research may reveal other relations between topographic properties and functional variation in the cerebellum.

Here we tested the UCT theory by analyzing in vivo structural and functional neuroimaging data in healthy humans. Future investigations might test the presence of similarities across motor, cognitive, and affective neuroimaging abnormalities in patients with isolated cerebellar injury. Neurophysiological investigations might also test the presence of similar physiological properties across distinct cerebellar regions engaged in distinct functions.

### Conclusion

The UCT theory has been an influential framework for understanding cerebellar function and dysfunction since its proposition more than two decades ago (Schmahmann 1991, 1996, 2000, 2010). Based on extrapolations from the observation of a uniform cerebellar cytoarchitecture (Ito 1993, Voogd & Glickstein 1998) and supported by additional clinical (Schmahmann 2000), behavioral (Guell et al. 2015b), and iTBS/EEG (Farzan et al. 2016) investigations, the central anatomical and physiological correlates of this theory have been elusive to direct testing. The modern neuroimaging and data analysis approaches in the present study allowed us to directly test and confirm the cerebellar form-function relation that UCT theory predicts. Our study provides a strong test of the idea that there are fundamentally different relations between microstructural variations and functional variations in the cerebellum versus the cerebral cortex. The absence of a microstructure-function relation in the cerebellum supports UCT theory that cerebellar computations are constant across functional domains.

## Acknowledgements

This work was supported in part by the MINDlink foundation (JS), la Caixa Banking Foundation (XG), and the MGH ECOR Tosteson - FMD Postdoctoral Fellowship Award (XG). Data were provided by the Human Connectome Project, WU-Minn Consortium (Principal Investigators: David Van Essen and Kamil Ugurbil; 1U54MH091657) funded by the 16 National Institutes of Health and Centers that support the Nation Institutes of Health Blueprint for Neuroscience Research; and by the McDonnell Center for Systems Neuroscience at Washington University.

